# Intra- and inter-chromosomal chromatin interactions mediate genetic effects on regulatory networks

**DOI:** 10.1101/171694

**Authors:** O. Delaneau, M. Zazhytska, C. Borel, C. Howald, S. Kumar, H. Ongen, K. Popadin, D. Marbach, G. Ambrosini, D. Bielser, D. Hacker, L. Romano-Palumbo, P. Ribaux, M. Wiederkehr, E. Falconnet, P. Bucher, S. Bergmann, S. E. Antonarakis, A. Reymond, E. T. Dermitzakis

## Abstract

Genome-wide studies on the genetic basis of gene expression and the structural properties of chromatin have considerably advanced our understanding of the function of the human genome. However, it remains unclear how structure relates to function and, in this work, we aim at bridging both by assembling a dataset that combines the activity of regulatory elements (e.g. enhancers and promoters), expression of genes and genetic variations of 317 individuals and across two cell types. We show that the regulatory activity is structured within 12,583 Cis Regulatory Domains (CRDs) that are cell type specific and highly reflective of the local (i.e. Topologically Associating Domains) and global (i.e. A/B nuclear compartments) nuclear organization of the chromatin. These CRDs essentially delimit the sets of active regulatory elements involved in the transcription of most genes, thereby capturing complex regulatory networks in which the effects of regulatory variants are propagated and combined to finally mediate expression Quantitative Trait Loci. Overall, our analysis reveals the complexity and specificity of cis and trans regulatory networks and their perturbation by genetic variation.

## Introduction

A decade of unbiased genome-wide association studies revealed that most of the genetic determinants of complex traits likely reside in non-coding regions of the genome (Maurano et al., 2012; Nica et al., 2010; Nicolae et al., 2010) and are genetic variants modulating gene expression; usually called expression Quantitative Trait Loci or eQTLs (Pickrell et al., 2010). As a consequence, large genome-wide collections of eQTLs across multiple Human populations (Lappalainen et al., 2013), conditions (Quach et al., 2016) and tissues (GTEx Consortium, 2015) have been collected and offer now the resources to systematically identify the genes and tissues involved in complex traits (Ongen et al., 2016). We do not understand precisely the downstream effects of eQTLs on the regulatory machinery of the cell, yet recent developments in high-throughput assays that capture biochemical changes (e.g. ChIP-seq (Johnson et al., 2007)), open regions (e.g. ATAC-seq (Buenrostro et al., 2013)) and conformation (e.g. Hi-C (Lieberman-Aiden et al., 2009)) at the chromatin level provide the tools to achieve this task. This has already given key insights into the molecular mechanisms underlying eQTLs function and the consensus picture that now emerges involves the binding perturbation of transcription factors which result in variable activity of the corresponding regulatory elements (Spitz and Furlong, 2012). These effects then propagate to distal regulatory elements and genes through chromatin interactions (Grubert et al., 2015; Kasowski et al., 2013; Kilpinen et al., 2013; McVicker et al., 2013; Waszak et al., 2015) whose possible range is delimited by chromatin contact domains (usually called TADs for Topologically Associating Domains) which represent the structural units of the three-dimensional nuclear organization (Lieberman-Aiden et al., 2009; Pombo and Dillon, 2015; Rao et al., 2014).

To further characterize gene regulatory networks and their perturbation by genetic variants, we extended our previous work (Waszak et al., 2015) and assayed a combination of three well-studied histone modifications (H3K27ac, H3K4me1 and H3K4me3), which are known to reliably capture enhancer and promoter activities, in 317 lymphoblastoid cell lines (LCLs) and 78 primary fibroblasts derived from unrelated European individuals (**methods 1-2**). We complemented this dataset by profiling gene expression and genotyping millions of variants in each sample. By leveraging the population-scale design of this study, we explore the interplay between genetic variants, regulatory activity and gene expression, and show how this captures interactions between regulatory elements, reflects the three-dimensional nuclear organization, varies across two cell types and propagates various forms of genetic effects on gene expression.

## Results

### 1. Inter individual variability of chromatin activity reveals Cis Regulatory Domains

As an initial step to comprehend the inter-individual variability of chromatin activity, we first integrated all chromatin assays into two matrices comprising 271,417 chromatin peaks quantified in 317 LCLs and 78 fibroblast lines from unrelated European individuals, respectively (**supplementary figures 1-2**) and then proceeded with chromatin Quantitative Trait Locus mapping (cQTL). To properly deconvolve biological from technical variability and therefore to maximize the power of cQTL discovery, we repeated the procedure multiple times across variable sets of covariates (**supplementary figures 3-4**; **methods 4-6**). Using the optimal configuration, we discovered 44,492 and 14,476 cQTLs for LCL and fibroblast, respectively, which constitutes to our knowledge the largest available collection of cQTLs. As expected, the variability of a substantial fraction of chromatin peaks results from nearby genetic variants (pi1=30.4%; see **supplementary methods D6** for a description of pi1) often located within open chromatin regions and exhibiting small allelic biases (i.e. same proportions of negative and positive effect sizes for variants located within chromatin peaks; **supplementary figures 5A-C**). Interestingly, the chromatin peaks under genetic control tend to cluster together (**supplementary figure 5D**) possibly revealing the coordination between nearby regulatory elements (Kilpinen et al., 2013; Waszak et al., 2015).

To further characterize this coordinated behavior, we systematically measured in LCLs the inter-individual correlation between nearby chromatin peaks (within 1Mb of each other) and found a strong signal of correlation (pi1=18.7%; **methods 7**) that rapidly decays with distance, slightly varies across ChIP-seq assay pairs, is driven by the most variable chromatin peaks (**supplementary figures 6A-C**) and importantly forms well-delimited domains that we call Cis Regulatory Domains (CRDs; **figure 1A-C**). We then looked at the fibroblast data to infer the cell type specificity of the significant correlations and found that long range correlations (e.g. > 100kb) are less shared between cell types than short range ones (e.g. < 100kb; **supplementary figure6D**); suggesting that they participate more to the regulatory program responsible for cell type specificity.

**Figure1:**
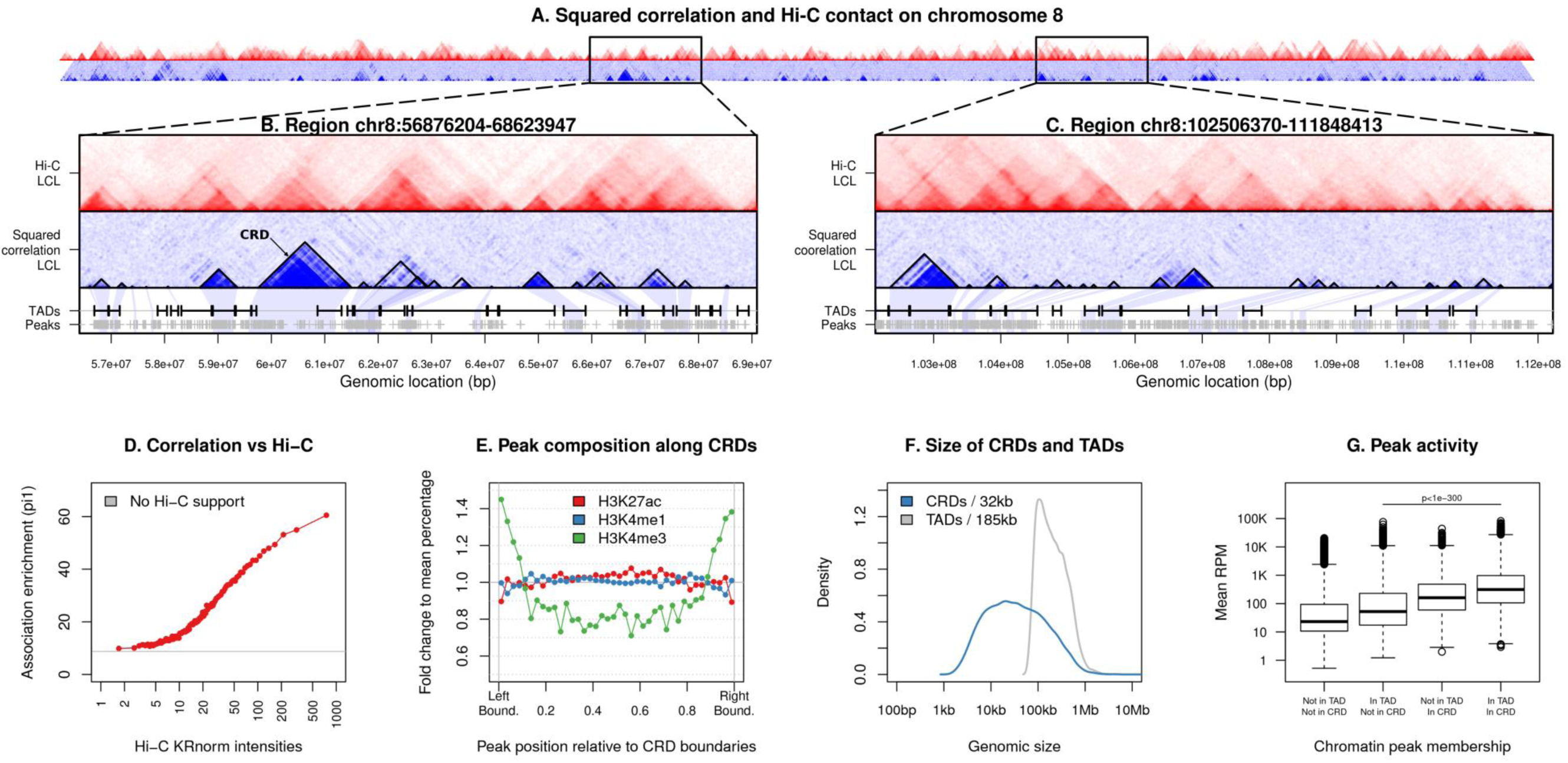
The *Cis* Regulatory Domains (CRDs). The squared correlations and Hi-C contact intensities (scaled between 0 and 1) between nearby chromatin peaks on chromosome 8 are shown in panel A with shades of blues and reds, respectively. Two examples containing 1,000 peaks are also shown in panels B and C together with (i) the CRDs (black triangles), (ii) the TADs (black segments) and (iii) the chromatin peak locations (grey crosses). The concordance between Hi-C and the correlation is shown in panel D: the association enrichment in the full LCL data was measured within 100 bins of increasing Hi-C contact intensities. The chromatin peak density as we move along CRDs is shown in panel E. The x-axis represents the position scaled between 0 and 1 of the chromatin peaks relative to the left and right boundaries of CRDs. The panel F compares the lengths of CRDs and TADs. And finally, the panel G gives the chromatin peak activity depending on their TAD and CRD memberships and expressed as Reads Per Millions.

We then searched for specific DNA properties that could underlie this correlation structure. To this aim, we first used publicly available deep Hi-C data for LCLs (Rao et al., 2014) to estimate the contact intensity between any pair of chromatin peaks (**methods 6**) and compared the resulting contact intensity measures to the correlation values. Overall, we find extremely good concordance between both measures (**figure 1D**). Then, we used a collection of protein binding sites derived for LCL as part of the ENCODE project (Encode Project Consortium, 2012) (**methods 3**) and found that the correlation decay is slower between binding sites of structural proteins (CTCF, RAD21, SMC3 and ZNF143; **supplementary figure6E**). Altogether, this shows that this correlation structure sufficiently captures the local three-dimensional conformation of chromatin.

A genome-wide call set of CRDs effectively delimits the proximal regulatory elements that likely interact together and therefore would help to decompose the genome into independent cis regulatory units. To assemble them, we designed a calling algorithm based on hierarchical clustering: chromatin peaks are iteratively grouped together into clusters based on the correlation levels they exhibit and the resulting binary tree is then cut at the level maximizing the total correlation mass captured (**methods 8**). This allowed us to discover 12,583 and 10,442 CRDs for LCL and fibroblast, respectively. In LCL, the CRDs are 32kb long on average, cover ^~^8% of the genome and span point associations between two distinct regulatory regions to complex association patterns that can encompass up to 80 distinct regulatory regions (**supplementary figures 7A-B**). By data subsampling, we assessed the discovery potential of this approach and find that 150 samples are enough to saturate the number of CRDs discovered while more samples are needed to better delimit the CRD content and boundaries (**supplementary figures 7C-D**). In terms of chromatin peak composition, we found that enhancers are the main component of CRDs (59.4% of H3K4me1 peaks), that they are uniformly distributed along them while promoters tend to locate at their boundaries (H3K4me3; **figure 1E**; **supplementary figure 7E**). This latter observation suggests that most of the CRDs we detected reflect how enhancers are brought in close proximity of promoters by chromatin looping.

### 2. CRDs delimit functionally active and variable TAD subdomains

Given the high concordance between Hi-C contacts and correlation data, we compared CRDs to a collection of TADs derived from a deep LCL Hi-C experiment that constitutes the most detailed definition of TADs so far (**methods 3**). CRDs are substantially smaller than TADs (32kb versus 185kb; **figure 1F**) even though they can reach the same size in some cases (up to 1Mb). When comparing the genomic regions defined by both CRDs and TADs, we find that 90% of the TADs overlap at least one CRD, 57% at least two and that CRDs tend to be fully encompassed within TAD boundaries (**supplementary figures 8A-B**). In addition, we also observed a higher density of CTCF binding sites at CRD boundaries similar to what is known for TADs (**supplementary figure 8C**) (Rao et al., 2014). All this constitutes accumulating evidences that CRDs actually correspond to TAD subdomains.

We then aimed at explaining why some of these TAD sub-domains correspond to CRDs while some others do not. To do so, we categorized the chromatin peaks into four categories depending on their location relative to both TADs and CRDs. First, this showed that chromatin peaks within CRDs are very likely to also fall within TADs (OR=1.75, fisher p-value < 1e-300); confirming the similarity between chromatin activity and structure. When focusing exclusively on TAD specific chromatin peaks and looking at their activity across all individuals, we find that those that are also members of CRDs exhibit much higher mean activity and variability than those that are not (**figure 1G**; **supplementary figure 8D**). Together, this suggests that TADs mix together inactive (potential) and active (realized) subdomains and we propose that active ones correspond to CRDs.

### 3. Inter-chromosomal correlation between CRDs reflects nuclear organization of DNA

By using fluorescence microscopy or Hi-C assays, multiple studies have characterized a high level organization of the chromatin; within chromosome territories (Meaburn and Misteli, 2007), A/B nuclear compartments (Lieberman-Aiden et al., 2009) and sub-compartments (Rao et al., 2014). Here, we used population variability of chromatin activity as perturbation to uncover the interactions underlying the global chromatin architecture. Specifically, we measured inter-individual correlations for all pairs of chromatin peaks located on distinct chromosomes (n=^~^3.4e10 tests) and found a substantial signal of association (pi1=8.1%; estimated by data sub-sampling). Given the difficulty to properly control for FDR in this massive multiple-testing setting, we first examined the data by considering as significant any pair of chromatin peaks with an association P-value below 1e-6. By doing so, we observed that some chromosomes preferentially associate with some others (**figure 2A**); notably chromosomes 16, 17, 19, 20 and 22 in line with the observation that these chromosomes tend to locate at the center of the nucleus thereby increasing their chance to interact with other chromosomes (Kaufmann et al., 2015). We also found that the per-chromosome association frequency (i.e. the fraction of tests per-chromosome with a p-value < 1e-6) correlates well with gene density (**figure 2E**) suggesting that gene rich chromosomes participate more to inter-chromosomal chromatin interactions (Kalhor et al., 2011). Consistent with this hypothesis, chromosomes 16, 17, 19, 20 and 22, are the richest in CpG islands (Lander et al., 2001). By visual inspection of the correlation structure for some pairs of chromosomes, we observed that strong correlations often concentrate at specific genomic locations that frequently involve CRDs (**figure 2B-C**). To validate these “hot spots” of correlation, we again used deep Hi-C data for LCL and found good concordance between the correlation structure and the inter-chromosomal contact intensities both locally (**supplementary figure 9**) and globally (**figure 2D**). Altogether, this suggests a global correlation structure between chromatin peaks above CRDs that reflects how chromatin is packaged into the nucleus.

**Figure2:**
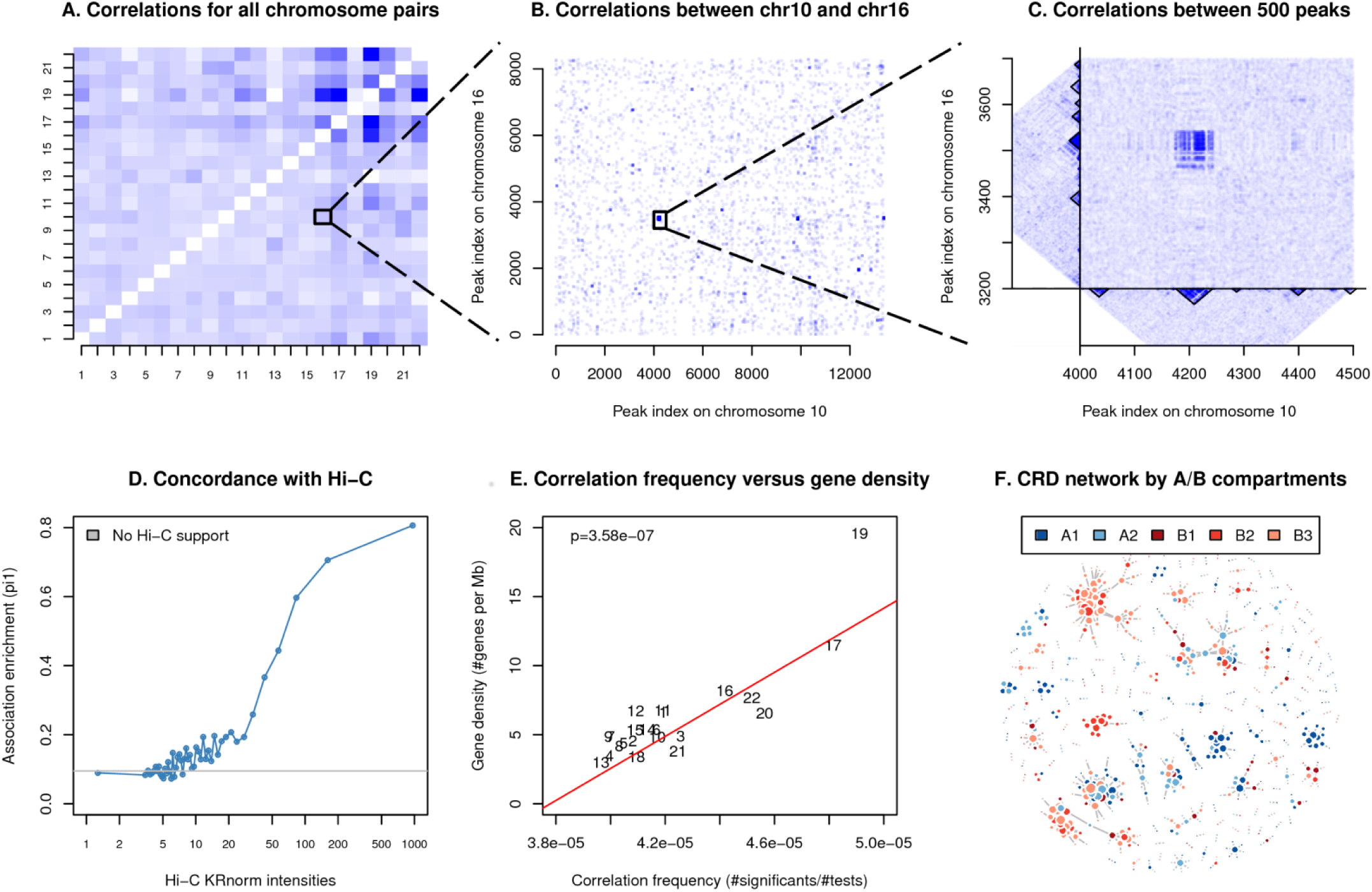
The inter-chromosomal interactions of chromatin. The three first panels show the correlation structure between chromosomes at three different zoom levels: (A) the frequency of correlations with a P-value < 1e-6 for all pairs of chromosomes, (B) all pairs of peaks between chromosome 10 and 16 with a P-value < 1e-4 and (C) a 500×500 peaks region exhibiting high correlation; all this in the context of CRDs and local correlation between chromatin peaks. The panel D gives the association enrichments of inter-chromosomal correlations within 50 bins of increasing Hi-C contact intensities. The panel E gives the association frequency as a function of the gene density on a per-chromosome basis. The panel F shows a network representation of the 1,000 strongest inter-chromosomal associations between CRDs colored by nuclear compartment membership.

To move from this global pattern to individual associations between regulatory elements while properly controlling for FDR, we next performed inter-chromosomal correlation analysis between CRDs. This requires quantifying CRD activity on a per-individual basis, a task that we performed using dimensionality reduction techniques to aggregate together the quantifications of multiple peaks of a CRD into a single quantification vector (**methods 8**). As a result, we reduced the number of tests by three orders of magnitude (from n=^~^3.4e10 tests to n=^~^7.5e7) and substantially increased the association signal (from pi1=8.1% between chromatin peaks to pi1=16.9% between CRDs; **supplementary figure 10A**). Thus, CRDs are commonly associated across chromosomes as illustrated by the 25,315 significant CRD-CRD associations detected at 1% FDR. Interestingly, bringing all these associations together into a network reveals the presence of hubs in which few CRDs play a central role and connect together multiple CRDs from distinct chromosomes (**figure 2F**; **supplementary figures 10B-C**). This led us to further refine our understanding of the global correlation structure of chromatin activity by annotating each CRD as belonging to one of the previously characterized nuclear compartments (A1, A2, B1, B2 or B3; **methods 3**) (Rao et al., 2014). From this, we made three observations: (i) CRDs from the same chromosomal compartment are more likely to be correlated, (ii) hubs usually bring together CRDs from the same chromosomal compartment and (iii) associations between CRDs from the same chromosomal compartment are more shared across cell types (**figure 2F**; **supplementary figures 10D-E**). All this shows that inter-chromosomal associations between CRDs are abundant in the genome, highly cell-type specific and form hubs with respect to how chromatin segregates into A/B nuclear compartments.

### 4. CRDs are fundamental functional units for gene regulation

The population scale design of this study offers a unique opportunity of building a genome-wide map interconnecting genes and CRDs at a resolution and a level of accuracy inaccessible to simple approaches based on co-localization or genomic distance. Indeed, we could directly assign genes to CRDs by assessing the significance of the correlation between their respective quantifications (**methods 9**). In practice, this approach allowed us to unravel a massive signal of associations: in LCL, the vast majority of the genes (pi1=82.4%) are associated with at least one CRD in cis (+/- 1Mb) which converts into 12,204 CRD-gene associations at 1% FDR (**figure 3A**; **supplementary figure 11A**). Often, these associations show relatively high connectivity degree; in 23.3% of the cases a gene is associated with multiple CRDs, while in 20.2% of the cases a CRD is associated with multiple genes (**figure 3B**). In terms of the linear position in the chromosome, we found that 46.8% of the associations connect genes to CRDs that encompassed their TSS (Transcription Start Sites), which also means that less than half of the associations are detectable through co-localization. Overall, the TSS of the associated genes tends to locate close to the CRD boundaries and usually exhibit a strong association signal when it falls within CRDs (**figure 3C**; **supplementary figure 11B**). In terms of histone mark composition, CRDs associated with genes are enriched for active marks (e.g. H3K27ac, H3K4me1 and H3K9ac), depleted for repressive (e.g. H3K27me3; **supplementary figures 11C-D**) and more often belong to the active nuclear compartment A (**figure 3D**). Altogether, this suggests that these associations bridge active CRDs to their target genes, thereby revealing local regulatory networks involved in the regulation of a vast majority of the genes.

**Figure3:**
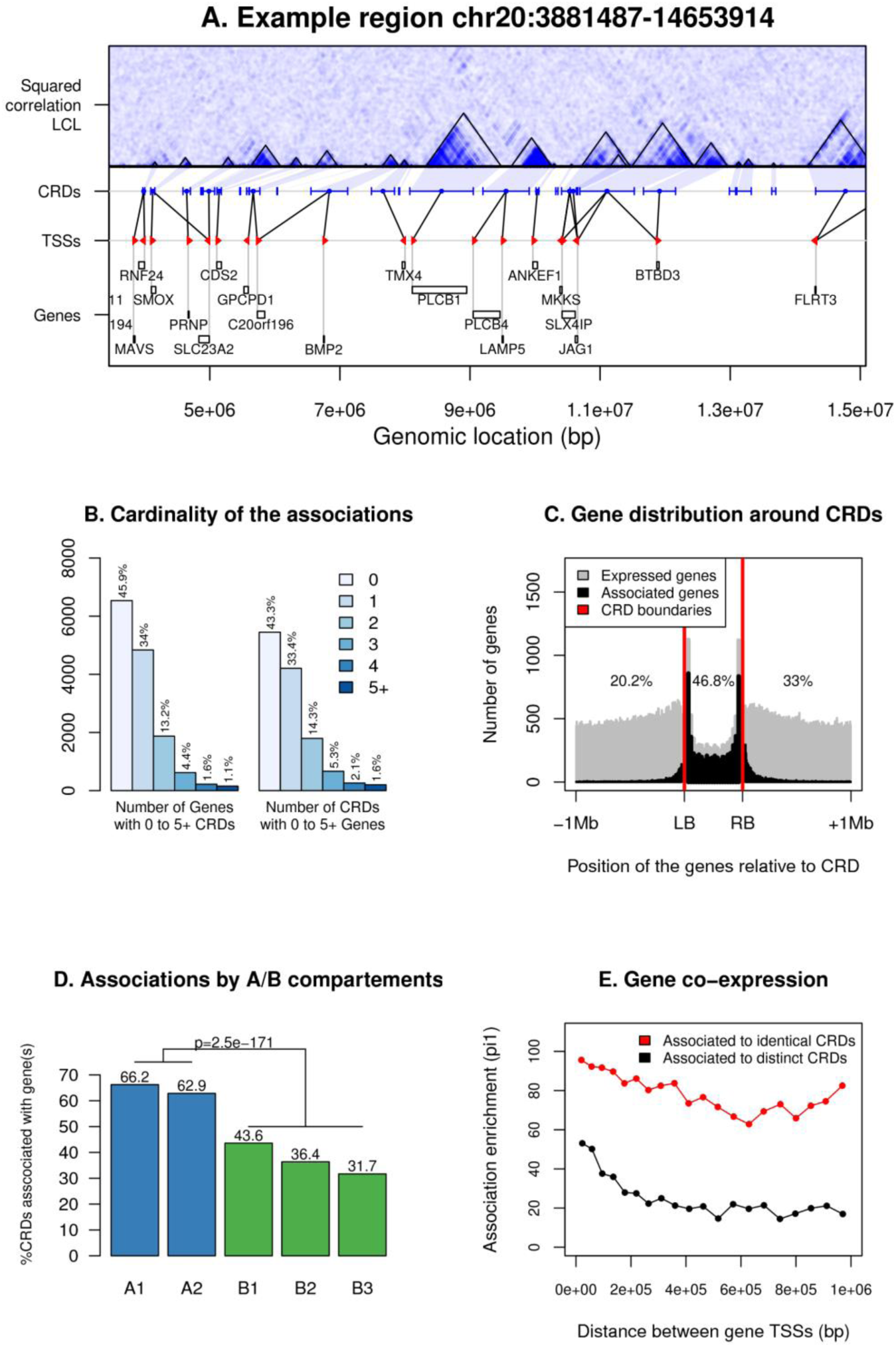
The regulatory function of CRDs. The first panel shows the associations between CRDs (blue segments) and genes (gene TSSs with red arrows and gene body with black boxes) within an example region of chromosome 20. Then, the panel B shows the connectivity between genes and CRDs while the panel C the distributions of tested genes (in grey) and significantly associated genes (in black) (i) within CRDs (relative locations; LB and RB correspond to left and right CRD boundaries, respectively) and (ii) within the genomic regions surrounding CRDs (absolute positions +/- 1 Mb). Panel (D) shows the percentage of CRDs being associated with genes within A/B nuclear compartments (significance comparing A to B is shown on top). The panel E gives the association enrichment in between genes associated with identical (in red) or distinct (in black) CRDs as a function of the distance between them.

The observation that genes and CRDs frequently show high connectivity and do not overlap led us to perform additional analyses. Notably, we first found that nearby genes linked to a single CRD are much more correlated with each other than those that are linked to distinct CRDs. Similarly, CRDs associated with the same gene are also highly correlated with each other (**figures 3E, supplementary figure 12A**). Distant gene-CRD associations (separated by from 10kb to 1Mb) occur within TAD boundaries more likely than what is expected by chance and exhibit more cell type specificity than short range ones (**supplementary figures 12B-C**). There is therefore another layer of coordination between nearby CRDs that is cell type specific and results in co-expression between nearby genes at the transcriptional level. This also illustrates the limitations of using domain representations alone to capture the hierarchical nature of regulatory interactions as additional layers of interactions are systematically missed.

### 5. CRDs are central mediators of genetic effects on gene expression

The relatively large sample size of the LCL data set (n=317) offers the required statistical power to analyze the genetic effects on CRDs. We therefore searched for Quantitative Trait Loci (QTLs) for both genes and CRDs. Concerning CRDs, we did not only quantify their activity as described before but also their structure by quantifying for each individual whether its data contributed positively or negatively to the overall correlation structure (**methods 8-9**). To summarize, we tested three molecular phenotypes for association with nearby genetic variants (**methods 4**), thereby resulting in three different kinds of QTL discoveries: at the gene expression level (eQTL), at the CRD activity level (aCRD-QTL) and at the CRD structure level (sCRD-QTL). In total, we discovered 6,157 aCRD-QTLs (pi1=68.5%), 7,658 eQTLs (pi1=59.2%) and 110 sCRD-QTLs (pi1=11.3%; **figures 4A-B**; **supplementary figure 13A**), which demonstrates that, similarly to genes, CRDs are under very strong genetic control. In terms of genomic distribution, all three types of QTLs exhibit the same pattern: they tend to become stronger and more frequent as we get closer to the genomic source of the target phenotype (**supplementary figure 13B**). However, in terms of minor allele frequency (MAF) spectrum, they show some differences: sCRD-QTLs involve almost exclusively genetic variants with a MAF below 20% while both eQTLs and aCRD-QTLs span the full MAF range (**figure 4C**); suggesting that the sCRD-QTLs are under intense selective pressure. This is also supported by the observation that sCRD-QTLs involve more often a loss of protein binding motifs compared to aCRD-QTLs and eQTLs (**supplementary methods D13**; **supplementary figure13C**). To summarize, this suggests two kinds of genetic variations affecting CRDs; the frequent case involving common genetic variants modulating the CRD activity through perturbation of protein binding affinity and the more deleterious case involving rare genetic variants that change the CRD structure by disrupting internal interactions.

**Figure4:**
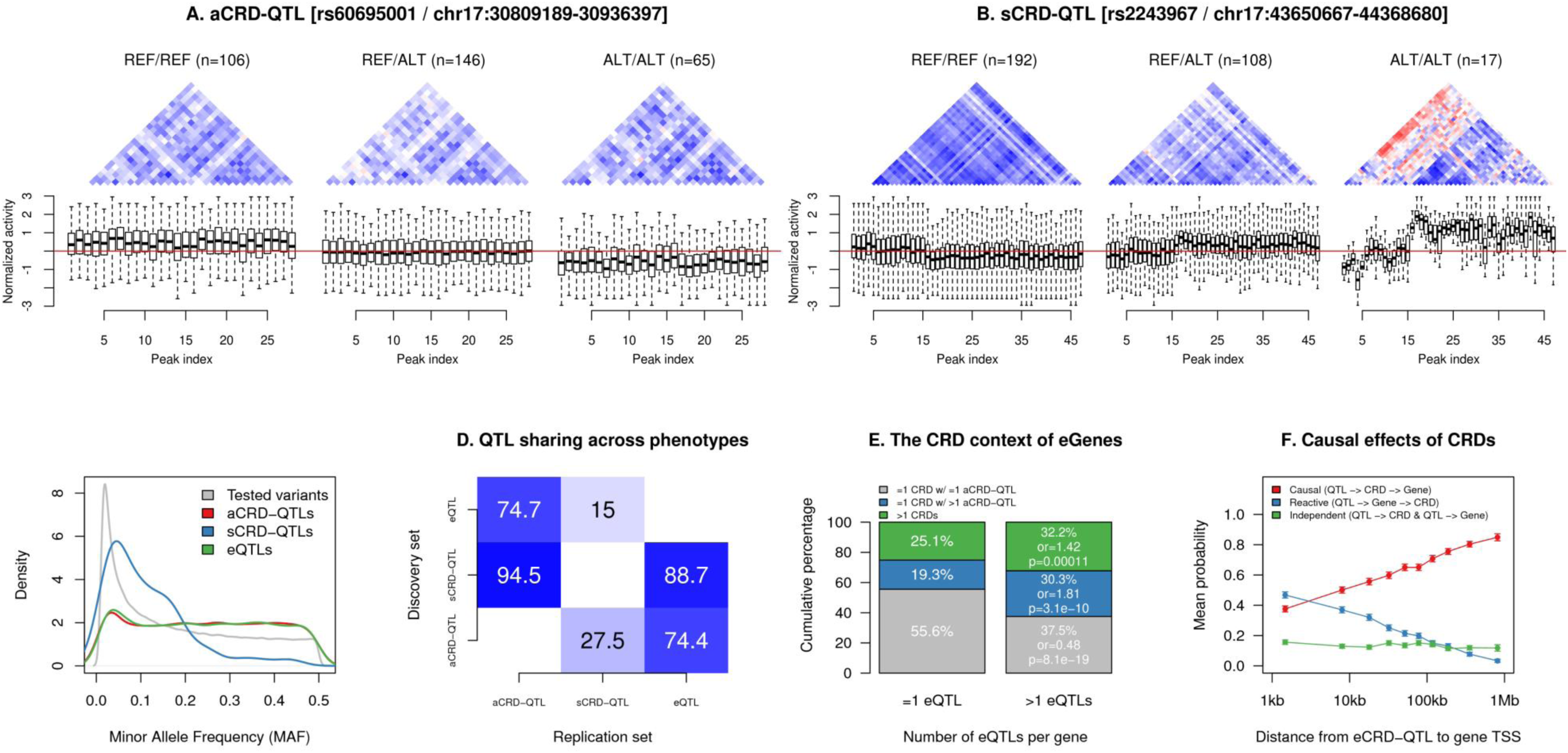
The genetic control of CRDs. The panels A and B show an example of aCRD-QJL (CRD-activity Q.TL) and sCRD-QJL (CRD-structure Q.TL), respectively. In each case, the correlation structure (positive and negative correlations in blue and red, respectively) and the normalized activity distribution (bottom panels) for all chromatin peaks contained in the CRD are shown stratified by the QTL genotype (REF/REF, REF/ALT and ALT/ALT). The numbers of individuals carrying each genotype are shown on top of each panel. The panel C shows the minor allele frequency spectrum for all types of QTLs while the panel D gives the association enrichments of QTLs across all molecular phenotypes (i.e. overlaps or degree of sharing). The panel E shows the respective proportions of genes with one eQTL or more being associated with (i) one CRD having one aCRD-QTL (in grey), (ii) one CRD with two or more aCRD-QTLs (in blue) and (iii) two or more CRDs (in green). The panel F gives the mean probability of each causal model as a function of distance between the eCRD-QTLs and the target genes. Only data for eCRD-QTLs falling within their target CRD is shown here.

As a natural step forward, we then assessed the extent to which these molecular QTLs overlap. In the context of our study, this task is greatly facilitated by the availability of a dense genome-wide map interconnecting genes and CRDs, which, in practice, allows testing for association QTLs of CRDs with relevant genes. Overall, we found extensive overlaps between all types of molecular QTLs (**figure 4D**). More specifically, we first observed that sCRD-QTLs are almost fully recapitulated by aCRD-QTLs (pi1=94.5%); indicating that perturbations of the CRD structure also result in activity variations. We then found that pi1=74.4% of the aCRD-QTLs affect gene expression, while pi1=74.7% of the eQTLs affect CRD activity. Thus, most of the molecular QTLs affect both CRD activity and gene expression and therefore likely tag the same causal variants. To identify the slight differences between both QTL sets, we compared their respective chances of overlapping functional annotations of the genomes (Encode Project Consortium, 2012; Whyte et al., 2013; Zarrei et al., 2015). The aCRD-QTLs locate more frequently in open chromatin regions, protein binding sites and enhancers, while eQTLs are enriched in promoters and transcribed regions (**supplementary figure13D**). Both types of QTL exhibit therefore different trade-offs in capturing distal/proximal genetic effects on gene expression.

We next examined the role of CRDs in mediating genetic effects on gene expression. First, we tried to look at how multiple eQTLs with independent effects on a given gene actually decompose into individual effects at the CRD level. We performed conditional analyses to supplement our previous QTL discoveries with additional independent ones (**supplementary methods D14**) and then compared the numbers of QTLs per gene and CRD by leveraging the gene-CRD associations. We have observed that genes with multiple eQTLs associate more often with (a) a single CRD having multiple aCRD-QTLs (OR=1.81) or (b) with more than 1 CRDs (OR=1.42) compared to genes with a single eQTL (**figure 4E**). This demonstrates that the joint effect of multiple eQTLs on a gene results from molecular interactions at two different levels: between regulatory elements within a single CRD or between distinct CRDs. Then, we asked how genetic effects actually flow through genes and CRDs in order to understand the causal relationships. A prerequisite of our analysis is to focus exclusively on genetic variants that affect both molecular phenotypes simultaneously. To detect these, we applied a Principal Component Analysis on each of the 12,204 gene-CRD associations and used the individual coordinates on PC1 as quantifications for QTL mapping. In total, we discovered 8,706 QTLs for gene-CRD pairs (called eCRD-QTLs; CRD-expression QTLs) and estimated that pi1=81.7% of these pairs mediate genetic effects. We then used Bayesian Network modeling to unravel causal relationships, an approach that has already been used for causal inference between genetic variants and molecular phenotypes (Gutierrez-Arcelus et al., 2013) (**methods 10**). In our study, this provided very high call rates in most of the cases (**supplementary figure 14A**); meaning that we were able to reliably assign a causal scenario for a vast majority of the 8,706 gene-CRD-QTL triplets under consideration. Overall, we find a clear signal that activity changes at CRDs are causal to changes in gene expression when they originate from eCRD-QTLs distal to genes while they are reactive for proximal eCRD-QTLs (**figure 4F**). Interestingly, this signal becomes even stronger when we only focused on eCRD-QTLs directly located within CRDs (**supplementary figure 14B**). This led us next to distinguish the functional elements that contribute to these two types of causal mechanisms (i.e. causal or reactive). We found that eCRD-QTLs involving a causal role for CRDs usually map in enhancers while those that involve a reactive role for CRDs map in promoters (**supplementary figure 14C**). Altogether, this shows that regulatory interactions within CRDs mediate the long range effects of enhancer related regulatory variations on gene expression while they actually reflect these perturbations in the case of promoter related regulatory variations.

The availability of four collections of molecular QTLs allows further functional comparisons. First, we can determine the degree to which those QTLs are shared across cell types. To do so, we quantified all the LCL features (CRDs, gene-CRD pairs and genes) in the 78 fibroblast samples and computed the P-values of association in fibroblasts of all significant LCL discoveries. We found that between pi1=27.3% and pi1=35.4% of the QTLs for CRDs are also significant in fibroblasts, a value which goes up to pi1=48% for eQTLs (**supplementary figure 14D**). Thus, QTLs for CRDs are more cell type specific than eQTLs similar to the previous observation that QTLs for CRDs capture more enhancer related effects. Second, we examined if our QTL collections are enriched for known GWAS hits (MacArthur et al., 2017) as it has already been shown for eQTLs (Maurano et al., 2012; Nica et al., 2010; Nicolae et al., 2010). We performed an enrichment analysis (**supplementary methods 17**) and found similar enrichment values across all types of QTLs, emphasizing their relevance in studying diseases, with a slight advantage for sCRD-QTLs as a likely result of their predicted deleterious effects (**supplementary figure 14E**).

We provide here two concrete examples how the dense genome-wide map interconnecting genes, CRDs and regulatory variations that we progressively assembled throughout this study could empower future association studies. As a first example, we used CRDs as a genome annotation to study the cumulative effect of multiple genetic variants. Specifically, we used whole genome and transcriptome sequencing data from the GEUVADIS study (**methods 3**) (Lappalainen et al., 2013) to assess if the accumulation of rare variants within a CRD is associated with the expression of nearby genes (i.e. rare-eQTL effects). To this aim, a ‘genetic burden’ was defined for each CRD by counting the number of minor alleles at rare variants (MAF<5%) falling into its regulatory regions and was then tested for association with the expression levels of nearby genes (**methods 11**). This amounts to a burden test at the regulatory level and showed that a non-negligible fraction of genes (pi1=10.4%) are affected by rare-eQTLs (**figures 5A-B**). At 5% FDR, this resulted in 33 CRDs whose baseline activity is perturbed by the accumulation of rare mutations with a level of expression of the downstream gene being decreased in 64% of the cases (**supplementary figure 15A**). This demonstrates the potential of our approach to explore the cumulative effects of multiple regulatory variations such as haplotypic effects. As a second example, we leveraged the inter-chromosomal associations between CRDs to discover trans-eQTLs likely acting through direct chromatin interactions and not diffuse factors (Bryois et al., 2014; Pierce et al., 2014). We implemented this by combining three different layers of associations: (i) aCRD-QTLs, (ii) CRD-CRD inter-chromosomal associations and (iii) gene-CRD associations. This defined 20,489 variant-gene pairs located on distinct chromosomes to be tested for association (**methods 12**), a task that we performed in six distinct datasets compiling data from two studies (the present study and EUROBATS; **methods 3**) and across five cell types (blood, fat, LCL, skin and fibroblasts). Overall, we found association enrichments at particularly high levels in the LCL datasets (**figure 5C**) potentially highlighting the cell type specificity of trans effects. We then extracted the 27 hits significant at 5% FDR in the largest LCL data set (EUROBATS LCL, n=765) and replicated 9 of them (p-value < 0.01 and same effect size direction) in our LCL dataset (n=317; **figure 5D**; **supplementary figure15B**). Almost none of these 27 hits are replicated in other cell types supporting the notion that they are real trans effects and not due to technical biases such as read mapping artifacts. By assembling all the relevant associations between CRDs, genes and trans-QTLs into networks, we find that all associations actually relate to two distinct variants (rs2980236 and rs7758352); the former variant affecting a total of 8 genes in trans, all of them connected by a CRD hub. Then, we enforced the directionality of the edges in the networks to match our mechanistic hypothesis that trans-eQTLs act through chromatin interactions and assessed whether this assumption is supported by the data using Bayesian networks (**supplementary methods 20**). Overall, the data provided extremely good support (**figure 5E**), thereby indicating that these 9 trans-eQTLs act through direct inter-chromosomal chromatin interactions.

**Figure5:**
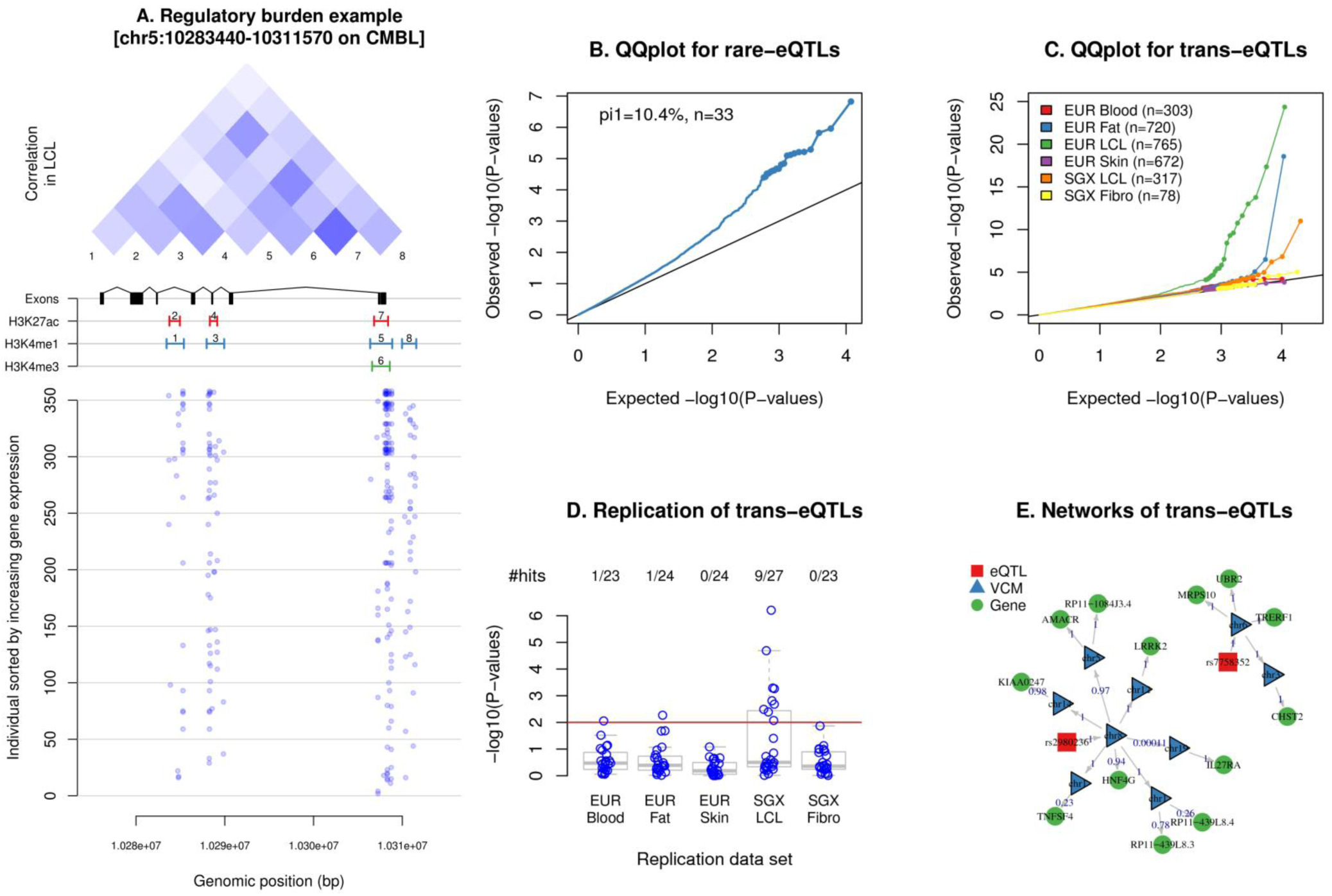
CRDs enhance association studies. The left panel (A) shows an example of how CRD can be used to perform a burden test of rare variants on gene expression. From top to bottom are shown (i) the pairwise correlations between peaks within the CRD, (ii) the positions of the various genomic features (peaks and gene) and (iii) the positions of the minor alleles at variants overlapping peaks for each of the 358 tested individual (sorted by increasing gene expression; low expression at the bottom). The QQplots in panel B and C show the association enrichments obtained when mapping *rare*-eQTLs (B; burden test) and *trans*-eQTLs in 6 distinct data sets (C). The panel D gives the replication P-values of the 27 significant hits found to be significant in EUROBATS LCL data. The panel E gives a network representation of the 9 replicated trans-eQTLs together with the relevant *cis* and *trans* CRDs and genes. The edge directions are set up given causal hypotheses and their robustness assessed using Bayesian networks: each weight ranges between 0 and 1; the latter meaning a direction of effect highly supported by the data.

## Discussion

In this study, we have analyzed genome-wide genotype, chromatin and gene expression data to reveal first- and higher-order regulatory interactions in the genome by using genetic variation as a perturbation. We illustrate the multiple benefits of integrating genome variation of multiple molecular assays to dissect the effects of regulatory variation onto gene expression at an unprecedented scale and resolution. We provide a very large set of chromatin QTLs that can form the basis of many specific analyses at the gene or region level in order to reveal genetic effects and interactions. We uncover the extensive interplay and organization of chromatin around genes and regulatory elements that ultimately form Cis Regulatory Domains (CRDs). These CRDs, while being embedded in Topologically Associating Domains (TADs), have higher complexity and degree of interaction at the sub-TAD level, capture local interactions with most genes and tend to be highly tissue specific. In addition, they are key factors involved in the co-regulation between nearby genes. However, they can also form a backbone for long-range direct interactions that result in regulation and co-regulation of genes at larger distances and on distinct chromosomes, revealing a level of organization that was previously suspected (Williams et al., 2010) but not documented beyond a few rare examples. To summarize, we find that genome variability in the context of regulatory activity and gene expression reflects the local and global nuclear organization of chromatin, thereby revealing complex networks of chromatin interactions. These regulatory networks are perturbed by genetic variations and serve as physical backbones to propagate and combine their effects on gene expression which can result in distal eQTL, independent eQTL, trans eQTL and rare eQTL effects. Overall, our work bridges discoveries made by genome-wide studies focusing on the structural properties of the chromatin together with those describing the genetic basis of chromatin activity and gene expression variability.

By drawing a dense and high resolution genome-wide map of the interplay between genetic variations, regulatory elements and gene expression across two cell types, we provide a unique reference resource to better interpret disease associated variants, a proof-of-principle that population-scale variability of molecular phenotypes is a powerful means to unravel the regulatory networks underlying gene expression and that these regulatory networks constitute informative priors to enhance future association studies. We also introduced major methodological advances related to the processing of multi-layered molecular phenotype data, notably for their integration through clustering and dimensionality reduction techniques and for causal inference through Bayesian networks. Future studies using such a population-scale approach should allow building on our work by including larger samples to capture even weaker effects, more cell types to ultimately build a cross-tissue catalog of regulatory networks, affected patients to characterize disease specific regulatory signature and other molecular assays to either rationalize the examination of active regulatory elements (e.g. ATAC-seq) or extend it to silencing effects (e.g. ChIP-seq for H3K27me3).

The consideration of genetic effects in local and global scale, as we have performed in this study, offers mechanistic insights that allow us to start assessing the true cellular impact of genetic variants individually as altogether and their impact not only at the gene level but also at the cellular level. This brings us closer to fully dissecting the properties and interactions of genetic variants defining organismal phenotypes via intra- and inter-cellular interactions.

## Author contributions

E.T.D., A.R., S.E.A., S.B. and P.B. designed and supervised the study and contributed to data interpretation. S.E.A. and C.B. provided the GenCord samples. D.H. cultured and processed the lymphoblastoid cell lines. C.B., P.R. and E.F. cultured and processed the fibroblasts. A.R. designed the ChIP experiments. M.Z. and M.W. executed the ChIP experiments. L.R-P. and D.B. prepared ChIP- and RNA-seq libraries and executed the sequencing. O.D. designed and executed the primary data analysis. C.H., G.A., H.O., K.P. and S.K contributed to data analysis. O.D. and E.T.D. performed the primary manuscript writing. A.R., S.E.A., S.B., D.M., C.B. and C.H. contributed to the manuscript writing.

## Data availability

All the raw sequence data (i.e. BAM files) generated in this study are deposited in the Array Express Archive (http://www.ebi.ac.uk/arrayexpress/ *[Available upon publication]*). All the processed data (molecular phenotype matrixes, collections of molecular QTLs and CRDs) are on a dedicated webpage (ftp://jungle.unige.ch/SGX *[Nicer website soon available]*). All the code implementing most computational methods used in this study has been packaged, documented and released on GitHub (https://github.com/odelaneau/clomics *[Better documentation soon available]*).

## Acknowledgments

This work was supported by grants from SystemX.ch, the Swiss Initiative in Systems Biology (SysGenetiX grant 3826), the Swiss National Science Foundation (SNSF) Sinergia grant CRSI33_130326 (ETD, AR), SNSF grants (31003A_160203 to AR and 163180 to SEA). The funders had no role in study design, data collection and analysis, decision to publish, or preparation of the manuscript.

## Methods

Hereafter is given a summary of all the experimental and computation methods used throughout this study. More details can be found in supplementary information and an overview of how all computational methods are interconnected is given in **supplementary figure 16**.

### 1. Sample collection (supplementary methods A)

We collected samples from three different sources: (i) 46 LCLs that we generated as part of a previous study (Waszak et al., 2015), (ii) 111 LCLs from the 1000 Genomes Project (1000 Genomes Project Consortium, 2015) and (iii) 160 LCLs and 78 fibroblasts from the GenCord collection (Dimas et al., 2009). Altogether, this gave a total of 317 LCLs and 78 fibroblasts all from individuals of European ancestry.

### 2. Experimental methods (supplementary methods B)

We cultured all cells for which no data was already available (271 LCLs and 78 fibroblasts) to perform both ChIP-seq and RNA-seq. Specifically, we performed chromatin immuno-precipitation for three different histone modifications: H3K27ac, H3K4me1 and H3K4me3 and extracted total RNA. We sequenced all resulting libraries on a HiSeq2000 machine. In addition, we also collected genotype data for all individuals by merging existing sequence data from the 1000 Genomes project (n=145) with data we generated using Illumina Human OMNI 2.5M SNP arrays (n=178).

### 3. External data (supplementary method C)

To analyze the data we generated, we used multiple external data sets: (i) contact maps, TAD definition and A/B nuclear compartments all derived for LCL from a deep Hi-C experiment (Rao et al., 2014), (ii) multiple functional annotations of the genome (binding sites for 50 transcription factors, 11 histone modifications, open chromatin regions and genome segmentation) derived for LCLs in the frame of the ENCODE project (Encode Project Consortium, 2012) and (iii) RNA-seq and genotype data for hundreds of European individuals from the GEUVADIS and EUROBATS projects (Buil et al., 2015; Lappalainen et al., 2013).

### 4. Genotype data preparation (supplementary methods D1)

All genotype data coming from SNP array have been merged together and filtered using standard procedures to remove low quality SNPs. The resulting genotype matrix was imputed from the 1000 Genomes phase 3 reference panel, poorly imputed variants removed and the rest combined with the genotype data coming from the 1000 Genomes sequencing data (1000 Genomes Project Consortium, 2015). This finally gave us genotype data for 323 individuals at 9,255,024 variants.

### 5. Molecular phenotype data preparation (supplementary methods D2-3)

All sequence data were mapped onto the human genome (hg19) using either BWA (Li and Durbin, 2009) for ChIP-seq data or GEM (Marco-Sola et al., 2012) for RNA-seq data. Gene expression has been quantified using QTLtools (Delaneau et al., 2017) with GENCODE v19 (Harrow et al., 2012) as reference and filtered to retain genes expressed in more than 90% of the samples. For the chromatin assays, we proceeded in two steps: we built a population scale BAM file for each assay by sub-sampling 1 million reads from 50 LCLs and 50 fibroblasts, then we performed peak calling on these using Homer v4.7 (Heinz et al., 2010) and quantified each sample in turn using the resulting peaks coordinates as reference genome annotations. We finally residualized all these molecular phenotypes for multiple covariates: sex, project of origin, genotyping platform, ancestry and technical variables (done by maximizing QTL discovery; **methods 6**). Of note, we merged the outcome of all chromatin assays together to obtain a total of four quantification matrices: (1) expression at 18,939 genes for 317 LCLs, (2) activity at 271,467 chromatin peaks for 317 LCLs, (3) expression at 18,068 genes for 78 fibroblasts and (4) activity at 271,467 chromatin peaks for 78 fibroblasts.

### 6. Molecular QTL mapping (supplementary methods D4)

We mapped molecular QTL for each molecular phenotype using (i) 1,000 permutations to correct for the number of genetic variants being tested in cis (+/- 1 Mb from the genomic feature boundaries) and (ii) the False Discovery Rate procedure implemented in the R/qvalue package (Storey and Tibshirani, 2003) to correct for the number of phenotypes being tested genome-wide. All these analyses were run using the QTLtools software package (Delaneau et al., 2017).

### 7. Building correlation and contact maps (supplementary methods D5-7)

We built correlation maps for the chromatin assays by systematically measuring inter-individual correlation (i.e. Pearson correlation coefficient) between any pair of chromatin peaks located on the same chromosome (intra-chromosomal) or on distinct chromosomes (inter-chromosomal). In multiple analyses, we measured the enrichment of small correlation P-values according to various parameters of interest by using the pi1 statistics of the R/qvalue package (Storey and Tibshirani, 2003). This pi1 statistics essentially gives an estimate of the number of statistical tests made under the alternative hypothesis of association. In addition, we built peak-centered Hi-C contact maps by interpolating the contact intensities between any pair of chromatin peaks from a deep Hi-C experiment made for LCL (Rao et al., 2014). Overall, this means that between any two chromatin peaks, we got two interaction measures: (a) the correlation between activity levels of the peaks and (b) the contact intensity measured by Hi-C.

### 8. Calling and quantifying CRDs (supplementary methods D8-10)

We called Cis Regulatory Domains (CRDs) from correlation data in two steps: (i) we performed hierarchical clustering on the chromatin data to get a binary tree for each chromosome that regroups the chromatin peaks depending on the correlation they exhibit and (ii) we cut the resulting binary tree at the level maximizing the total correlation mass captured by using three empirical criteria (i.e. overall, edge and distal correlations) which gave us clusters of highly correlated chromatin peaks (i.e. the actual CRDs). Once the CRDs were called, we quantified them on a per sample basis at two levels: (i) we measured their overall activity by simply taking the averaged peak quantification and (ii) we measured their structure by assessing the contribution of each individual onto the mean correlation within the CRD.

### 9. Mapping genes and QTLs for CRDs (supplementary methods D11-12)

Once the CRDs quantified, we searched for associations with nearby genes and genetic variants using the permutation procedure implemented in the QTLtools software package (Delaneau et al., 2017). This allowed us to efficiently correct for the fact that multiple genetic variants, genes and CRDs are tested genome wide. Of note, we only searched for associations within 1Mb of the CRD boundaries. For the genes, we used as genomic coordinates the position of their TSSs and only considered protein coding genes or lincRNAs.

In some cases, we complemented this step by conditional analysis as implemented in the forward-backward scan of QTLtools (Delaneau et al., 2017).

### 10. Inferring causality (supplementary methods D15-16)

To infer the causal relationships between QTLs, CRDs and genes, we proceeded in two steps. First, we only focused on QTLs affecting both CRDs and genes. To get those, we quantified each gene-CRD pair using PCA and mapped QTLs using the coordinates on PC1 as quantifications. As a byproduct, this gave us QTL-CRD-gene triplets onto which we performed Bayesian Network (BN) analysis to get the posterior probabilities for the three possible causal models we considered in this study: (i) the causal model with QTL > CRD > gene, (ii) the reactive model QTL > gene > CRD and (iii) the independent model with QTL > CRD and QTL > gene. This has been implemented using the R/bnlearn package (Scutari, 2010).

### 11. CRD based burden test (supplementary methods D18)

We designed a burden test at the regulatory level by leveraging CRDs. This test relies on two steps: (i) we measured the burden of rare mutations (minor allele at variant with a MAF < 5%) for each CRD and (ii) we then tested this burden with expression at relevant genes. By relevant genes, we mean genes that have been previously found to be associated with the CRD of interest. We performed this analysis on a dataset in which we have whole genome and transcriptome data for 358 individuals (1000 Genomes Project Consortium, 2015; Lappalainen et al., 2013).

### 12. CRD driven trans-eQTL mapping (supplementary methods D19-20)

We built a map of gene-variant pairs located on distinct chromosome to be tested for association by leveraging the inter-chromosomal CRD-CRD associations we discovered in the frame of this study. We assembled together (i) gene-CRD associations (cis links), (ii) CRD-CRD associations (trans links) and (iii) QTL-CRD associations (cis links) which effectively forms 20,489 gene-variant pairs to be tested in *trans.* Then, instead of performing the billions of tests required by standard trans-eQTL analysis (millions of variants times thousands of genes), we only focused on the 20,489 pairs this approach helped us to assemble. We performed these tests in multiple data sets (e.g. EUROBATS (Buil et al., 2015)) and extracted the significant hits we obtained at 5% FDR.

